# Task-irrelevant phase but not contrast variability unlocks generalization in visual perceptual learning

**DOI:** 10.1101/2024.02.01.578442

**Authors:** Beyza Akkoyunlu, Caspar M. Schwiedrzik

## Abstract

Performance on visual tasks can be improved by practice, a process called visual perceptual learning. However, learning-induced performance improvements are often limited to the specific stimuli and visual field locations used during training. Recent research has shown that variability along task-irrelevant stimulus dimensions during training can reduce this specificity. This has been related to higher stages of visual processing that harbor neurons which are invariant to the task-irrelevant dimension. Here, we test whether task-irrelevant trial-by-trial variability in two visual features for which invariances arise at different stages of processing, contrast and spatial phase, results in different degrees of generalization in space in an orientation discrimination task. We find that randomizing spatial phase results in complete generalization of learning to a new spatial location, contrary to randomizing contrast. Our results thus suggest that the neural population undergoing plasticity in visual perceptual learning is determined by the training task, which, in turn, affects generalization. This lends further support to the hypothesis that task-irrelevant variability is an independent factor in determining the specificity of perceptual learning.

## Introduction

Our visual environment is ever-changing. In turn, our brain adapts to this challenge by maintaining the ability to learn throughout life. This capacity also extends to our visual sense: through experience, we can mould and even expand our perceptual abilities (Dosher & Lu, 2020). Yet, how we learn from diverse experiences remains a lingering question.

In the laboratory setting, we stimulate learning in the visual system by employing visual perceptual learning (VPL) paradigms, e.g., training participants on orientation discrimination tasks. While VPL results in profound and long-lasting performance increases, these learning-induced gains are usually highly specific to the training parameters (Lu & Dosher, 2022). For instance, even if only the location or spatial frequency of the stimuli are changed after training, learning effects disappear (Fiorentini & Berardi, 1980; Schoups et al., 1995). This characteristic of VPL is scientifically intriguing, but limits the practical use of VPL, e.g., in vision restoration therapies.

Because generalization is often considered the ultimate goal of learning, recent research has asked the question whether it is possible to develop training regimes that overcome the characteristic specificity of VPL. One factor that has been shown to lead to generalizable learning effects in many other domains such as motor, language, or category learning is *variability* (Douvis, 2005; Perry et al., 2010; Vukatana et al., 2015). Variability may promote learning at a more abstract level and thus prevent “overfitting”, i.e., learning effects that are extremely specific to the training material (Raviv et al., 2022). Indeed, our own recent research has shown that systematically inducing variability in a task-irrelevant dimension can lead to more generalized learning effects even in visual orientation discrimination (Manenti et al., 2023), a domain that had so far been considered to yield only extremely specific VPL effects (Schoups et al., 1995; Shiu & Pashler, 1992). Although the results from this new training strategy answer one question, they raise another: does all variability lead to generalization?

VPL likely involves multiple plasticity mechanisms that occur throughout the brain (Maniglia & Seitz, 2018; Watanabe & Sasaki, 2015). To explain whether learning is specific or generalizes, theories often suggest that this depends on the stage of processing at which learning occurs in visual cortex. *Specificity* may arise when early stages of processing are involved, such as the primary visual cortex (V1), where neurons respond with high specificity to a narrow range of stimulus features, e.g., a particular orientation or spatial location (Fahle, 2005). Hence, if learning changes the tuning properties of these neurons, their readout, and/or how they are targeted by top-down processes, highly specific learning effects may ensue. On the other hand, *generalization* has been hypothesized to involve higher-level areas in visual cortex, where neurons respond to a broader range of features and wider regions of the visual field (Ahissar & Hochstein, 2004). *Variability* is thought to tap into the so-called invariance properties of these higher-order neurons: invariant representations consistently respond to task-relevant features irrespective of variability in other, task-irrelevant features. Lower brain areas could only achieve this by distributing plasticity over a large number of neurons co-tuned to the task-relevant and the task-irrelevant feature. Hence, relying on invariant neurons could provide task-relevant information in face of variability while simultaneously limiting the number of neurons or readout weights that need to undergo plasticity (Manenti et al., 2023). Alternatively, plasticity could always involve the lowest level of processing at which the relevant feature dimension is differentially encoded (Karni & Bertini, 1997).

To contrast these two hypotheses, we trained participants on an orientation discrimination task, which is known to lead to highly specific learning effects (Schoups et al., 1995; Shiu & Pashler, 1992). During training, we introduced trial-by-trial variability in one of two task-irrelevant dimensions, the spatial phase or the contrast of the stimuli, respectively. We opted for these dimensions because they constitute fundamental features for visual processing (Gladilin & Eils, 2015; Shapley, 1985), yet, above threshold, they are not essential for the orientation discrimination task. Furthermore, *invariance* to these two features arises at different levels of processing. In particular, phase invariance appears later in the visual processing hierarchy than contrast invariance: for instance, contrast invariance arises already in simple cells, whereas phase invariance is first seen in V1 complex cells (Hubel & Wiesel, 1962) and fully emerges in V2 (Cloherty & Ibbotson, 2015). As neurons at higher stages of processing typically respond to larger regions of the visual field (i.e., they have larger receptive fields), we hypothesized that plasticity involving such neurons would result in less spatial specificity. Alternatively, given that both contrast and spatial phase are not essential to orientation processing above threshold, both randomization types could result in a comparable degree of spatial specificity if plasticity always involved the lowest stage of processing where the task-relevant feature is encoded. We thus tested whether participants trained with phase randomized stimuli would achieve more transfer to untrained spatial locations than participants trained with contrast-randomized stimuli.

To preview, we find that phase and contrast randomization lead to differential spatial transfer after orientation discrimination learning. In particular, participants trained with phase randomized stimuli show generalization of learning effects in space that extend further than the learning effects of subjects trained with contrast randomized stimuli. This supports the idea that variability induces plasticity involving neurons higher in the visual processing hierarchy and suggests that the specific feature in which variability occurs can determine at which stage of processing this plasticity occurs.

## Materials and methods

### Participants

Forty healthy volunteers (18 female, 4 left-handed, mean age 25 years, SD 2.66 years) participated in the experiment. All participants had normal or corrected-to-normal vision, reported no neurological or psychiatric diseases, and gave written informed consent before participation. Subjects were assigned to one of four experimental groups: phase randomized upper visual field (UVF) training (n=9) or contrast randomized UVF training (n=9), phase randomized lower visual field (LVF) training (n=11) or contrast randomized LVF training (n=12). Participants received 12€/hour compensation, and to ensure proper motivation, they received a 2€ bonus each day if they improved their orientation discrimination threshold by at least 10% compared to the previous day. All procedures were in accordance with the principles put forward by the Declaration of Helsinki and approved by the Ethics Committee of the University Medical Center Göttingen (protocol number 29/8/17).

### Stimuli and procedure

Subjects were trained with a two-alternative forced choice (2AFC) orientation discrimination task using Gabor gratings (Figure 1A). Each subject participated in a pre- and a post-training threshold measurement, respectively, and four training sessions. Each session was performed on a separate day, and participants had to complete all sessions of the experiment within ten days. On average, subjects took 7.26 days (SD 1.2 days) to complete the study.

**Figure 1.**
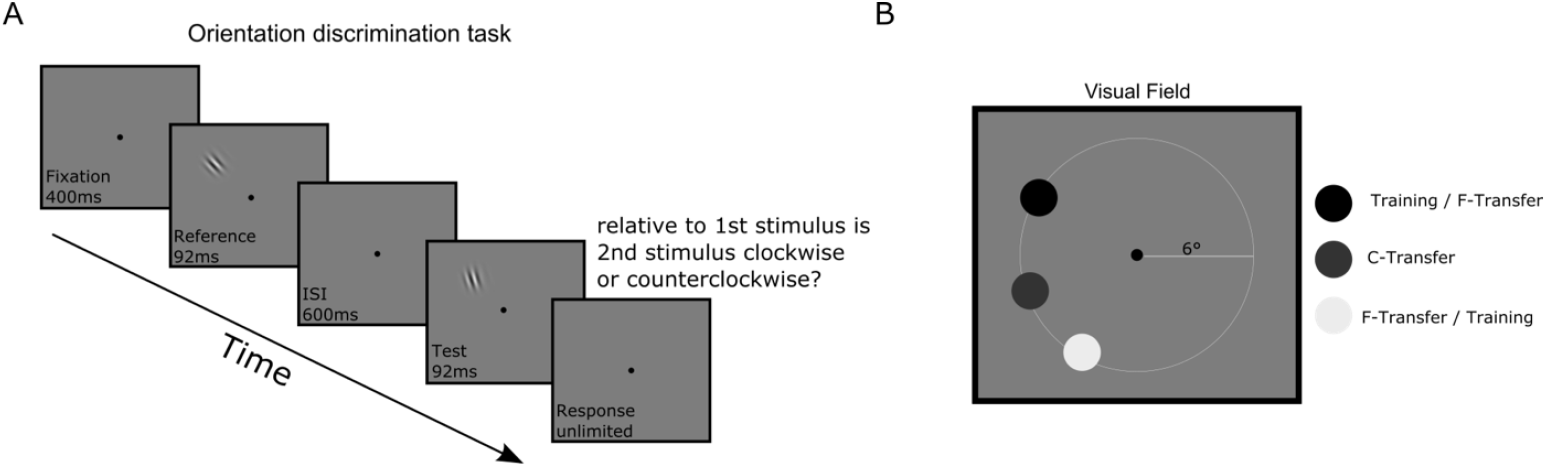
Task design. **A)** Each trial started with a 400 ms fixation period. Participants were then presented with the 45° reference grating (92 ms), followed by a 600 ms inter-stimulus interval, and then the test grating (92 ms). Participants were instructed to respond after the test grating disappeared. If participants broke fixation during the trial, the trial was aborted and repeated immediately. **B)** Training and transfer locations. All participants’ thresholds were collected at the respective training location and two transfer location, namely, close (C-Transfer) and far (F-Transfer) transfer locations. Training and F-Transfer locations were switched for the groups who trained in the upper visual field and lower visual field, respectively.

The stimuli were presented on a gamma-corrected LCD monitor (ViewPixx EEG, resolution 1920 × 1080 px, refresh rate 120 Hz) using Psychtoolbox (Brainard, 1997) running in Matlab (R2018B, The Mathworks). A circular aperture was placed on top of the screen to cover the monitor*’*s corners in order to prevent participants from using it as a reference while determining the orientation of the gratings. Participants sat at a distance of 65 cm from the screen and performed the task using a response box (Millikey SR-5). During the experiment, eye movements were recorded using a video-based eye tracker (SR Research Eyelink 1000+). Data were collected from the left eye at 1000 Hz.

Our experimental stimuli and procedure closely followed a study by Xiao and colleagues (2008). Gabor gratings (size 2 dva, spatial frequency 2.5 cpd, standard deviation of the Gaussian 0.28 deg) were placed at 6° eccentricity in the upper or lower left visual field, respectively, on a grey background (52.42 cd/m^2^ luminance). For the phase randomized group, the gratings were presented at a constant 45% Michelson contrast, and the spatial phase of the stimulus was randomized on each trial between 0° and 180°. In the contrast randomized group, spatial phase of the stimulus was kept constant at 0°, and contrast was randomized between 33% and 66% Michelson contrast. The fixation dot had a size of 0.23 dva.

In the threshold measurement sessions, participants performed three blocks of the 2AFC orientation discrimination task at each of three locations (Figure 1B). Participants started each trial with fixating for 400 ms, then a reference Gabor grating was presented for 92 ms. After a 600 ms inter-stimulus interval, the test grating was presented for 92 ms. Our stimulus timing (92 ms), taken from Xiao and colleagues (2008), falls squarely into the range of previously used stimulus durations, which vary between 33 ms (Lu & Dosher, 2004) and 500 ms (Tan et al., 2019) in the VPL literature. The reference grating had an orientation of 45° and the orientation of the test grating was determined with a 3 down – 1 up staircase procedure. Except for the orientation, reference and test gratings had identical parameters. Participants were instructed to report the orientation of the test grating in comparison to the reference grating (clockwise or counterclockwise). They were further instructed to respond as accurately and quickly as possible (but no limit on the responses time was imposed). If fixation was broken during the fixation period of the trial (2° tolerance window), the trial was aborted and immediately repeated. No feedback was provided.

Every block consisted of two interleaved staircases, one starting from a 1° offset and the other starting from a 3° offset from the 45° reference with 0.05 log step size. The blocks ended after 10 reversals or when 200 trials were completed (in a subset of session, more reversals were collected because of a coding issue). The thresholds were measured at three isoeccentric locations (Figure 1B), namely, training, close transfer (C-Transfer) and far transfer (F-Transfer). The distance between training – C-Transfer and C-Transfer – F-Transfer locations was 5°, whereas the distance between training – F-Transfer location was 11.5°. The order of testing at the three locations during the threshold measurement sessions was counterbalanced between participants and kept identical for pre and post training measurements.

Subjects were trained at one of two locations (upper or lower visual field) and with one of two types of task-irrelevant variability (spatial phase or contrast). In each training session, participants performed five blocks of 200 trials of the 2AFC task, i.e., 1000 trials per training day, for a total of 4000 training trials. The task design was identical to the threshold measurements, but participants now received feedback: error feedback as a high-pitch sound through headphones (Sennheiser HDA 280) after a mistake in a given trial; in addition, upon finishing the block, the percentage of correct trials was displayed on the screen to provide subjects with easily understandable performance feedback. Before each training session, participants were informed whether they had earned the financial performance bonus for the preceding training session in order to ensure continuous motivation.

### Analyses

One participant had to be excluded because he/she fell asleep during the experiment. Eleven participants were excluded because they did not learn (7 subjects did not learn at all and 4 subjects had a LI of 20% or worse; the same results were obtained using an exclusion criterion of a LI of 30% or worse). There was no statistically significant difference between number of participants excluded from each experimental group (Fisher exact test, p = 0.490, odds ratio = 0.201). The final n was 29 (14 female, 2 left-handed, mean age 24.5, SD 2.31).

All statistical analyses were conducted in R (version 4.2.2; R Core Team, 2022) on macOS Monterey 12.0.1.

Each subject performed three threshold blocks at each retinal location pre and post learning. To calculate the threshold of each block, the first four reversals of the staircase were discarded, and the mean of the following six reversals was taken. Blocks with an insufficient number of reversals were discarded. To determine how learning affected orientation discrimination thresholds at the training and transfer locations, we fitted a linear mixed effects model to the block-wise threshold data from each subject using the lme4 package (version 1.1-34) (Bates et al., 2015). Data were aligned and rank transformed (Higgins & Tashtoush, 1994) using the ARTool package (version 0.11.1) (Wobbrock et al., 2011), to satisfy distributional assumptions. In the linear mixed effects model, we predicted thresholds with the factors *time* (2 levels, pre/post), *randomization* (2 levels, phase/contrast), *location* (3 levels, training/C-transfer/F-transfer) and *training location* (2 levels, upper/lower visual field) as fixed effects, and participants as random intercept, as in Equation 1:

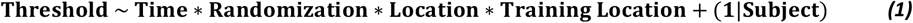

To obtain p-values, we performed a repeated measures analyses of variance (rmANOVA) of the model terms and their interactions using Wald’s F-test and degrees of freedom determined following Kenward and Roger (1997). All fixed effect variables are coded with center coding. Posthoc tests were performed on estimated marginal means of the aligned and rank transformed data (Elkin et al., 2021) using the emmeans package (version 1.8.8) (Lenth, 2023).

For quantifying learning, we also calculated the Learning Index (LI) (Xiao et al., 2008), as in Equation 2:

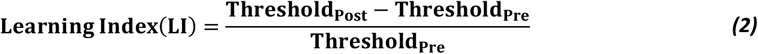

LIs were used to test whether transfer to the C-Transfer location correlated with transfer to the F-Transfer location across subjects; a one-sided Fisher’s z-test was used to compare the correlation coefficients between groups via the cocor package (version 1.1.4) (Diedenhofen & Musch, 2015).

To determine effect sizes, we calculated partial η^2^ for rmANOVAs, Cohen’s d for t-tests, and Cohen’s q for comparing correlation coefficients using the effectsize package (version 0.8.6) (Ben-Shachar et al., 2020). Plots were prepared using the ggpubr (version 0.6.0) (Kassambara, 2023) and ggplot2 (version 3.4.4) (Wickham, 2016) packages.

### Eye movements

Trials were aborted if subjects broke fixation during the fixation phase of the trial. Because trial-to-trial variability in eye position relative to the stimuli could induce additional phase variability beyond the variability induced by our experimental stimuli, we quantified fixation variability as the standard deviation of the average distance of the gaze location to the fixation point during the stimulus presentation across trials. In addition, we also quantified residual miniature eye movements, i.e., microsaccades (MS), which have previously been implicated in VPL (Hung & Carrasco, 2023). To this end, we first identified blinks in the data using the Eyelink parsing algorithm and excluded these timepoints as well as 100 ms before and 150 ms after the blink from analyses. Raw gaze position was converted to dva. Data were segmented into trials (200 ms before the reference stimulus presentation until 350ms after the offset of the target stimulus). We then detected MS using a velocity-based algorithm (Engbert & Kliegl, 2003). This algorithm calculates 2D velocities per trial and identifies MS candidates when the velocity surpasses 6 SD above the average velocity of the trial for a minimum of 6 ms. From these, MS candidates with less than 1° amplitude and a minimum 10 ms gap between MS were selected to prevent false positives (Kapoula & Robinson, 1986).

To calculate the MS rate per s, we calculated the average number of MS per time point (1 ms) across all trials in each session and multiplied these values by the sampling rate (1,000 Hz). To calculate MS percentage following Hung & Carrasco (2023), we divided the number of MS made in a session by the number of valid trials (trials which do not contain a blink) for each session and each participant. This was done for the time window of the reference stimulus, the target stimulus, as well as the whole trial.

## Results

We first examined whether there were any pre-existing differences between the contrast and phase randomization groups in terms of their pretraining orientation discrimination thresholds. This was not the case: at none of the three locations, there were statistically significant differences in pre-training thresholds between the contrast and phase randomized groups (Welch’s two-sample t-test, Training location t(25.214) = 0.702, p = 0.489, Cohen’s d = 0.28; C-Transfer t(25.868) = 0.011, p = 0.99913, Cohen’s d = 0.000; F-Transfer t(26.967) = 0.9348, p = 0.3582, Cohen’s d = 0.36). Hence, there was no evidence for pre-existing differences in orientation discrimination between the groups.

Upon training, both randomization groups improved in discriminating orientations at the training location: the group trained with phase randomization improved thresholds on average by 1.53 deg, and the group trained with contrast randomization improved on average by 1.40 deg. Learning in all groups was well characterized by a power law function (see Supplementary Figure S1), as has previously been found in some (Dosher & Lu, 2005) but not all studies (Cochrane & Green, 2021; Dosher & Lu, 2007) (see Supplementary Discussion).

We then assessed whether learning and transfer differed between the groups considering all three locations (Figure 2). Comparing learning (i.e., the pre-post difference in thresholds) at the training location, we found no statistically significant difference between the phase randomized and the contrast randomized group (t(469) = 1.062, p = 0.2888, Cohen’s d = 0.10). This suggests that despite different kinds of variability, training was similarly effective at this location. Transfer of learning to the C-Transfer location (again in terms of the pre-post difference in threshold) also did not reveal a statistically significant difference between the two groups (t(469) = 0.936, p = 0.3495, Cohen’s d = 0.09). This suggests that training transferred similarly to a close by location with both randomization manipulations. However, there was significantly more transfer to the F-Transfer location in the phase randomized group than the contrast randomized group (t(469) = 2.910, p = 0.0038, Cohen’s d = 0.27): while thresholds improved on average only by 1.31 deg in the contrast randomized group, the average improvement was 1.99 deg in the phase randomized group. This suggests that training effects generalize more broadly in space with phase than with contrast variability during training. This differential effect was also supported by a significant time × location × randomization interaction (F(2,469.031) = 3.317, p = 0.037, η^*2*^_p_ = 0.01, Supplementary Table 1 and Supplementary Figure S2). Overall, these results support the hypothesis that training with phase variability involves neurons with larger receptive fields than training with contrast variability.

**Figure 2.**
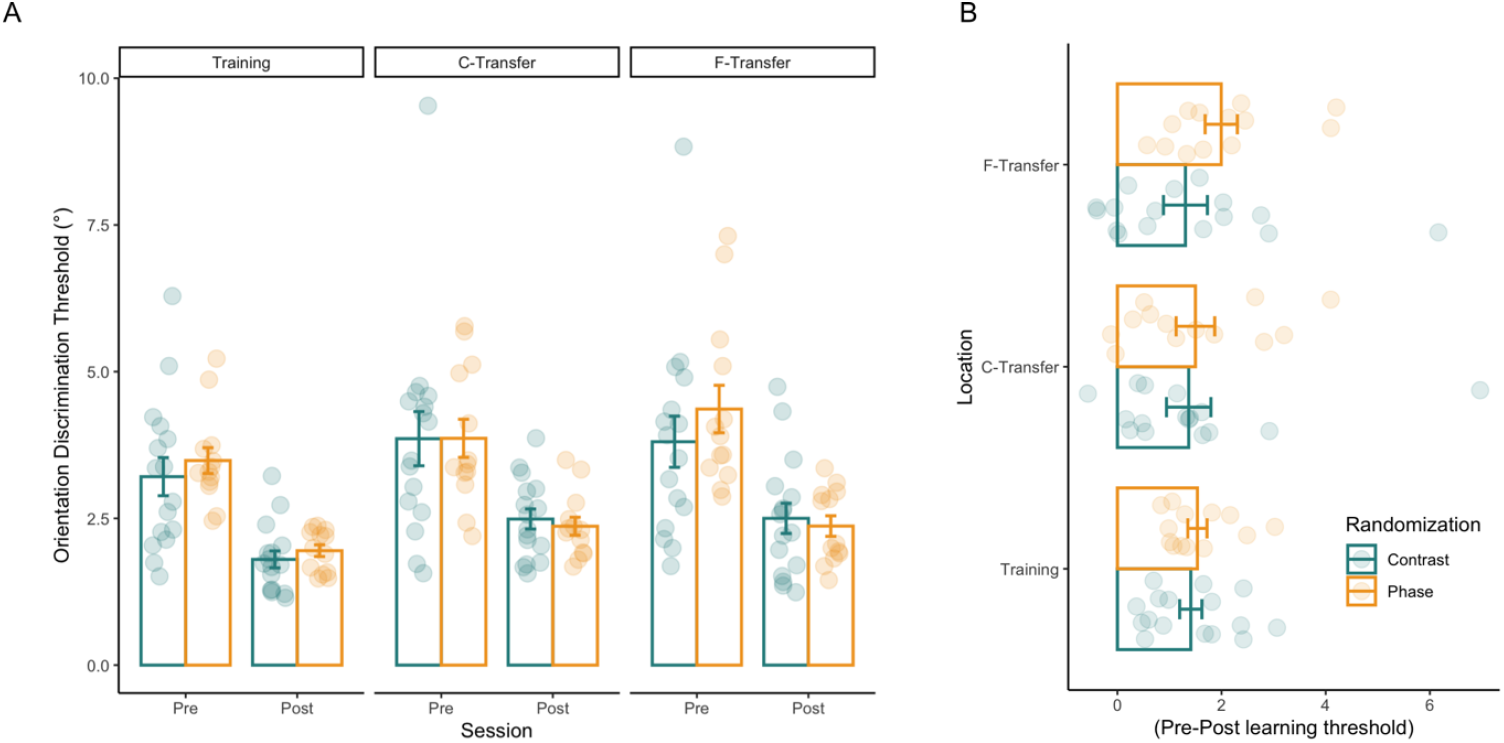
Pre and post learning thresholds at the training and transfer locations. **A)** Pre and post learning thresholds at all locations for phase (orange) and contrast (green) randomization. Each dot represents a subject’s mean threshold at the respective location and time point. Bars show the mean thresholds over subjects in each group and location. At the training location, there was no statistically significant difference between the phase randomized and the contrast randomized group (t(469) = 1.062, p = 0.2888, Cohen’s d = 0.10). Transfer of learning to the C-Transfer location also did not reveal a statistically significant difference between the two groups (t(469) = 0.936, p = 0.3495, Cohen’s d = 0.09). However, there was significantly more transfer to the F-Transfer location in the phase randomized group than the contrast randomized group (t(469) = 2.910, p = 0.0038, Cohen’s d = 0.27). **B)** Change in the thresholds with learning. Change in the threshold calculated per subject, by subtracting mean post-learning threshold from mean pre-learning threshold. In both panels, error bars represent the standard error of the mean.

Another prediction that can be derived from the hypothesis that phase randomization and contrast randomization rely on differently sized receptive fields is that learning effects should correlate between spatial locations if both locations fall into the same receptive fields, whereas they should not correlate if neurons with small, non-overlapping receptive fields are involved (Kondat et al., 2023). We thus investigated the correlation of LIs at the two transfer locations for both randomization groups (Figure 3). We find that LIs at the C-Transfer and F-Transfer locations correlate highly in the phase randomization group (two-sided Pearson*’*s correlation, r = 0.71, t(11) = 3.39, p = 0.006). In contrast, there was no statistically significant correlation of LIs between the two transfer locations in the contrast randomization group (two-sided Pearson*’*s correlation, r = 0.09, t(14) = 0.373, p = 0.71). Furthermore, the correlation in LIs was significantly larger in the phase than in the contrast randomized group (z = 1.8964, p = 0.0290, Cohen’s q = 0.79, one-sided). This provides further evidence that the two groups rely on different neural populations for learning orientation discrimination.

**Figure 3.**
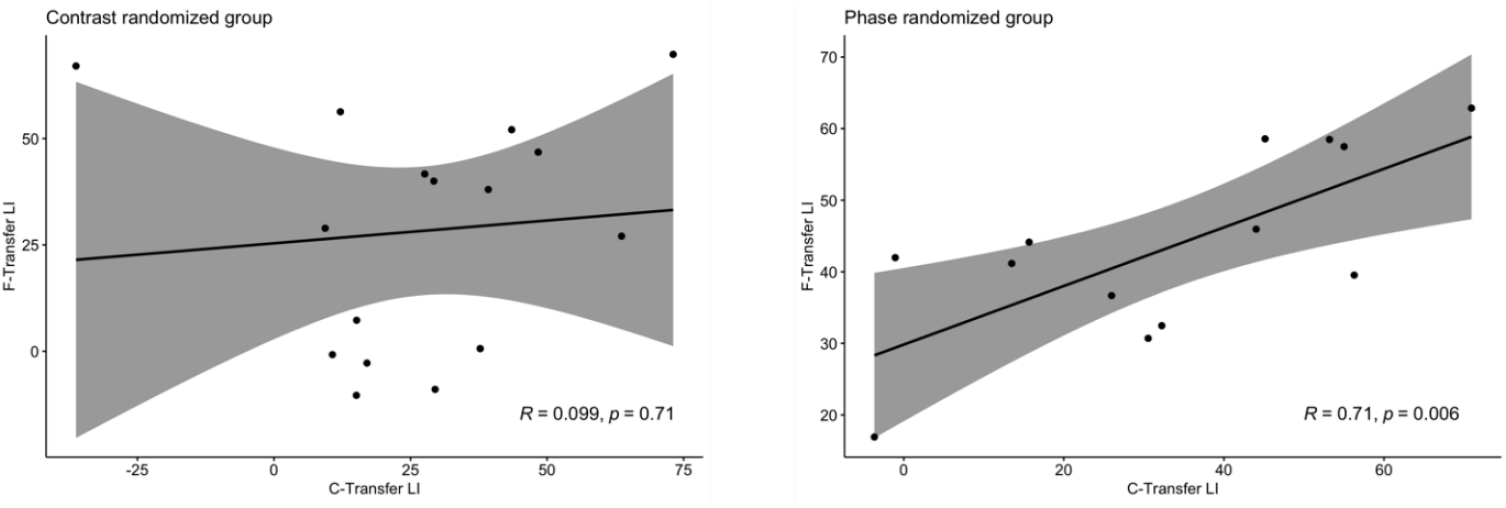
Correlation of LIs between the two transfer locations. In the contrast randomized group, LIs at the C-Transfer and F-Transfer locations are uncorrelated (r = 0.09, p = 0.71, two-sided). This result also holds when excluding a subject with an extreme LI of −36% at the C-Transfer location. In contrast, LIs are correlated in the phase randomized group (r = 0.71, p = 0.006, two-sided). The correlation in LIs was significantly larger in the phase than in the contrast randomized group (z = 1.8964, p = 0.0290, Cohen’s q = 0.79, one-sided). Shaded areas represent the 95% confidence intervals. Note that different scaling is used in each panel to ease comparison.

### Fixation variability results

Inspection of eye movements revealed that subjects generally fixated well within the fixation window, but that residual eye movements did occur that could in principle create additional variability during training. For example, in the contrast-randomized group, subjects showed variability in gaze position from trial to trial that may have led to a change in the fixated stimulus period across trials. However, retinal phase did not change in over a third of the trials and the amount of eye movement induced change was overall not comparable to the amount of stimulus-induced variability in the phase-randomized group. To statistically evaluate whether there were any differences in fixation variability between our experimental groups contributing to the differential effect, we performed an ANOVA, testing the effects of randomization and training location. The results do not reveal any effect of randomization (F(1,25) = 1.78, p = 0.19, *η_p_^2^* = 0.07), training location (F(1,25) = 3.93, p = 0.06, *η_p_^2^* = 0.14), or interaction of training location and randomization (F(1,25) = 0.70, p = 0.41, *η_p_^2^* = 0.03).

We then tested whether higher fixation variability leads to better generalization across locations. To this end, we correlated LIs with fixation variability. Both C-Transfer and F-Transfer location LIs did not correlate significantly with fixation variability (Figure 4; all p > 0.058). Together, this suggests that eye movement induced variability in spatial phase does not explain differences in the generalization of orientation discrimination learning effects between the groups.

**Figure 4.**
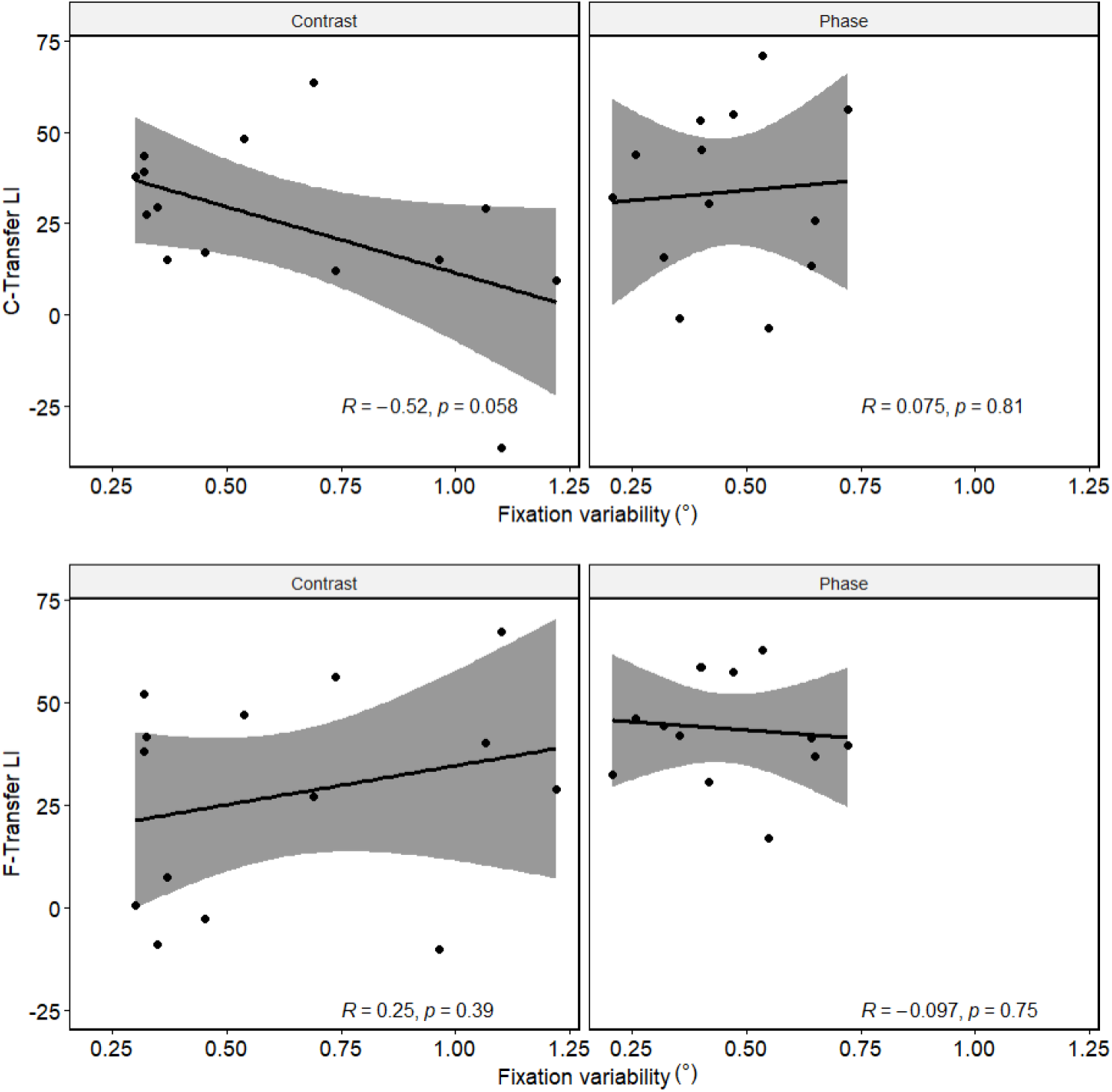
Correlation of fixation variability (in visual degrees) and transfer LIs. Both C-Transfer (upper section) and F-Transfer (lower section) LIs are not significantly correlated with fixation variability and these results hold for both randomization types (all p > 0.058). Shaded areas represent 95% confidence intervals.

### Microsaccade results

MS velocities and amplitudes were highly correlated (Pearson’s correlation, r=0.69, t(43618)=199.76, p<2.2×10^−16^), i.e., they followed the so-called “main-sequence”. After having established that we had correctly detected MS, we examined how MS varied with our experimental factors. We first tested whether MS percentage during target stimulus presentation changed with session, randomization and/or training location. MS percentage during the target stimulus presentation did not change with session (F(5,123.150)=0.63, p = 0.67, η_*p*_^*2*^=0.02), randomization (F(1,24.954)=0.45, p=0.50, η_*p*_^*2*^=0.02), or training location (F(1,24.954)=1.12, p=0.29, η_*p*_^*2*^=0.04), and there were no statistically significant interactions (all p>0.05). Similarly, MS percentage during reference stimulus presentation did not vary depending on session, randomization, or training location (session F(5,123.047)=0.41, p=0.83, η_*p*_^*2*^ =0.02; F(1,24.985)=0.11, p=0.73, η_*p*_^*2*^=0.00; training location F(1,24.985)=0.24, p=0.62, η_*p*_^*2*^=0.00; all interaction effects p>0.05). When considering the entire trial, MS percentages changed slightly with session (F(5,123.051)=3.07, p=0.01, η_*p*_^*2*^=0.11), but we did not find any statistically significant effect of randomization (F(1,24.984)=2.06, p=0.16, η_*p*_^*2*^=0.08), training location (F(1,24.984)=1.88, p=0.18, η_*p*_^*2*^=0.07), or interaction (all p>0.05). Hence, the different stimulation parameters did not lead to statistically appreciable differences in the occurrence of MS.

We then assessed whether MS were related to the differential generalization effects we found in our experimental groups. To this end, we correlated LIs at the two transfer locations, respectively, with MS percentage. These analyses did not reveal any evidence for MS leading to more generalizable learning effects (Figure 5, all p>0.08).

**Figure 5.**
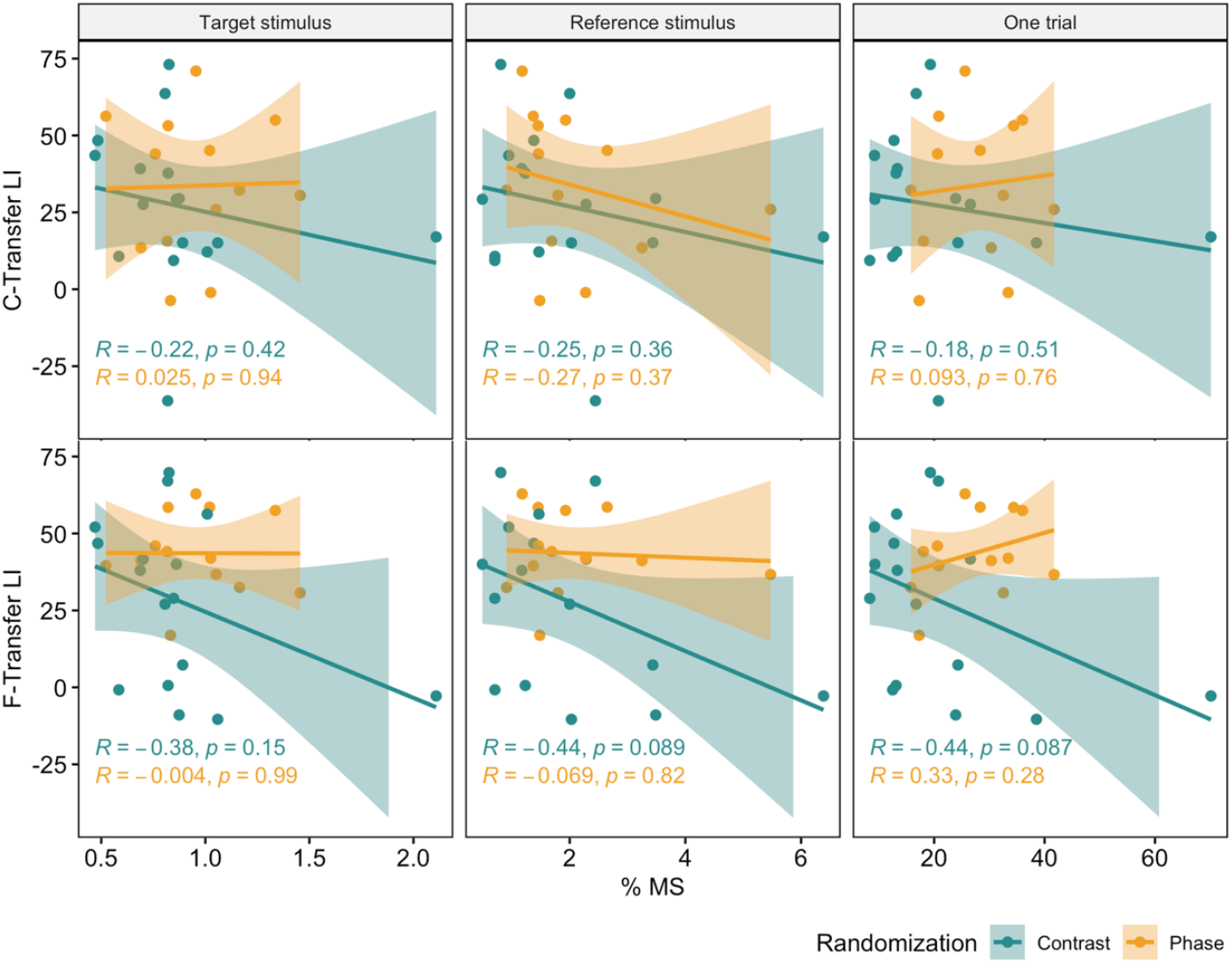
Correlations between microsaccade (MS) percentage and LIs. The upper row shows correlations at C-Transfer and the lower row at F-Transfer during different periods: during the target stimulus, reference stimulus, and for the entire trial (which includes ISI and both stimuli). There were no significant correlations in both randomization groups in any of the trial phases tested (all p>0.08). Shaded areas represent 95% confidence intervals.

## Discussion

We find that variability in spatial phase during training leads to more generalization of VPL in space than variability in contrast. Moreover, participants trained with variable spatial phase transfer learning similarly across two transfer locations, while subjects trained with variable contrast do not. Taken together, these two findings support the hypothesis that the two forms of variability tap into neurons with differently large receptive fields, and in turn, that the type of variability introduced during training can determine the stage of visual processing involved in VPL.

It has been suggested on the basis of behavioral data and simulations that when our visual system is confronted with variability in a task-irrelevant dimension, it can rely on neurons that are invariant to this feature and thus subsume this variability (Manenti et al., 2023); in the brain, such neurons typically reside at a higher stage of visual processing and have large receptive fields. Alternatively, the visual system can make use of neurons at earlier stages of processing that are conjunctively tuned to the task-relevant and the task-irrelevant features of the stimuli. Because these neurons typically have spatially restricted receptive fields, they cannot, by themselves, enable generalization across the visual field. Importantly, different invariances arise at different levels of processing. Hence, creating variability in different task-irrelevant features of a stimulus should result in different patterns of generalization. This is what we find for the case of spatial phase and contrast variability.

Relevant to our experiments, it is known that contrast invariance arises earlier in the visual processing hierarchy than phase invariance: contrast invariance gradually increase along the visual ventral stream (Avidan et al., 2002), but is already seen at the level of simple cells in V1 (Hubel & Wiesel, 1962). The earliest stage at which phase invariance arises is on the level of complex cells in V1 (Hubel & Wiesel, 1962). Both simple and complex cells are orientation tuned. According to a classical model of orientation tuning in V1, simple cells feed into complex cells (Hubel & Wiesel, 1962), and there is empirical evidence that supports this view (e.g., Alonso & Martinez, 1998). This principle also underlies hierarchical models of visual processing such as HMAX (Serre & Riesenhuber, 2004). Furthermore, simple cells are predominantly found in the granular input layer of V1, whereas complex cells are found in supra- and infragranular output layers (Hubel & Wiesel, 1968). Hence, complex cells reside on a higher level of processing than simple cells already in V1. Full phase invariance only arises in complex cells in V2 (Cloherty & Ibbotson, 2015), and thus later than contrast invariance.

Importantly, V1 complex cells have larger receptive fields than simple cells (Schiller et al., 1976), especially at higher eccentricities (Wilson & Sherman, 1976), and fully phase-invariant complex cells in V2 have larger receptive fields than contrast-invariant simple cells in V1 (Gattass et al., 1981). Hence, the earliest stages of processing at which differential spatial transfer effects between contrast- and phase-randomized training could hypothetically occur would be between simple and complex cells in V1, and/or between V1 and V2. Of note, simple and complex cells are not only found in V1, but also at higher stages of processing such as V2 (Levitt et al., 1994) and V4 (Desimone & Schein, 1987). Hence, the principle that phase invariance arises at a higher stage of processing than contrast invariance holds irrespective of where exactly plasticity occurs. Which stage of processing is actually involved and in which form cannot be resolved through behavioral experiments alone but will require future neurophysiological and/or neuroimaging studies.

We note that the spatial generalization that we observe is substantial relative to the typical receptive field size in early visual areas, including V4. Yet, neurophysiological recordings in macaque monkeys have shown that orientation tuning can be found not only in early visual areas, where it is typically studied, but that high-level visual cortical areas in the temporal lobe with large receptive fields (Desimone & Gross, 1979) contain orientation-tuned neurons (Gross et al., 1972; Vogels & Orban, 1994) that are causally relevant for orientation discrimination (Adab & Vogels, 2016). These could possibly support the large distances between the different locations in our experiments with their larger receptive fields. Unfortunately, at least to our knowledge, the difference in receptive field size between simple and complex cells has not been mapped for higher visual areas and the distinction between simple and complex cells has not been made for orientation tuned neurons that have been found beyond area V4 in the macaque monkey. Neurophysiological and/or neuroimaging experiments would be necessary to achieve this. Nevertheless, the distinction between simple and complex cells as different stages of processing seems to hold irrespective of these unknowns, as does the emergence of full phase invariance at a higher stage of processing than contrast invariance. Our results are thus in line with the hypothesis that the dimension in which task-irrelevant variability occurs determines the stage of visual processing involved in VPL, reflected by varying degrees of generalization.

Previous VPL studies have used phase or contrast randomization not as an experimental variable, but to assure that subjects are not using spatial cues to solve orientation discrimination tasks (in the case of spatial phase) and/or to reduce adaptation. Some of these studies have reported spatial generalization effects (e.g., Zhang et al., 2010). In contrast, a study by Sowden and colleagues (2002) reported training effects that were extremely specific to the spatial frequency, retinal location and even eye of origin, consistent with learning effects in the granular input layer of V1 where neurons which such tuning properties reside. Importantly, this study used a training procedure with no variability in which only a single grating was presented for several thousand trials. Notably, a study investigating the neural basis of orientation discrimination learning in macaque monkey V1 used phase randomization and reported learning effects only outside the granular input layer (Schoups et al., 2001), which would be compatible with an outsized involvement of complex cells. Overall, this suggests that by using variability during training, previous studies may have coincidentally engaged neurons at higher stages of processing and with larger receptive fields.

Apart from variability, spatial generalization of VPL also occurs when adaptation is reduced or eliminated (Harris et al., 2012). While spatial transfer to the C-Transfer location we found in both randomization groups may be explained by a reduction of adaptation with variability, the differential transfer effects at the F-Transfer location and the differential correlation between learning effects at the two transfer locations in the different groups are not easily accounted for by adaptation. Previous studies have also shown that task difficulty affects generalization (Ahissar & Hochstein, 2004). However, as pretraining thresholds did not differ significantly between contrast and phase randomization, this factor also cannot easily explain our differential transfer results. Hence, variability may contribute to generalization in VPL beyond the well-established factors task difficulty and adaptation. On the other hand, studies using contrast randomization during orientation discrimination learning in the context of double training (Xiao et al., 2008) and manipulation of attention (Donovan & Carrasco, 2018) have reported sizable spatial transfer effects. These may rely on more intricate neural mechanisms such as top-down influences. Hence, variability is not the only factor that enables spatial transfer after perceptual learning, but there are alternative and/or additional paths and mechanisms of achieving generalization.

While participants were instructed to direct their gaze to the fixation point and this was enforced by eye-tracking, deviations of gaze position within the fixation window (2°) can introduce additional variability of retinal phase beyond the experimentally induced stimulus variability. This is because even small displacements in fixation can lead to phase variability in the receptive field of a given neuron, including in the contrast randomized group. We investigated whether this potential source of variability affected spatial generalization of learning, yet, we did not find that fixation variability within the fixation window differentiates the different experimental groups or leads to higher generalization. Similarly, small, involuntary eye movements in the form of MS could in principle constitute an additional form of variability beyond the variability in stimulus features that we induced in our experiments. Interestingly, it has previously been proposed that such miniature eye movements increase stimulus discriminability (Ahissar & Arieli, 2012; Hennig et al., 2002; Melloni et al., 2009; Rucci et al., 2007). Furthermore, MS have been found to change in frequency over the time course of perceptual learning (Hung & Carrasco, 2023). However, we did not find any differences between the training regimes that would support a specific role of MS in the differential spatial generalization patterns that we observed. We note that recordings in macaque V1 have found no differential effect of eye movements on simple versus complex cells (Kagan et al., 2002). It has previously been proposed that lateral connections in V1 implement a form of spatial invariance that counteracts the possibly detrimental effects of positional uncertainty brought about by miniature eye movements during perceptual learning (Otto et al., 2006; Zhaoping et al., 2003). Alternatively or additionally, physiological (Ringach, 2002; Victor & Purpura, 1998) and behavioral (Field & Nachmias, 1984; Huang et al., 2006) data indicate that phase tuning is relatively broad. This would suggest that substantial phase variability (such as the phase variability in our experimental condition, which exceeded eye movement induced variability) is needed to tap into phase-invariant stages of processing. Ultimately, future experiments using gaze-contingent displays and/or stimuli with different spatial frequencies less prone to the effects of eye movements will be needed to differentiate the effects of stimulus-versus eye movement-induced variability on spatial generalization that we could not fully dissociate in our study.

While our study exemplifies the differential effects of different kinds of task-irrelevant variability, it is important to emphasize that our study only investigated a narrow range of stimulus parameters in each group and had a limited sample size. E.g., as mentioned above, a narrower range of phase variability might result in higher specificity. Similarly, performance is known to vary across the visual field (Abrams et al., 2012) which may lead to pre-training differences, which may only be uncovered with larger sample sizes. Predicting outcomes of VPL with a much wider range of variability or other features and/or locations becomes more challenging given multi-level nature of VPL (Maniglia & Seitz, 2018). E.g., extending the range of contrast variability might cause interactions with other factors such as task difficulty (Talluri et al., 2015), or perceived variability (Zaman et al., 2021) that could affect behavioral outcomes. Hence, further systematic studies with improved methods, such as non-random group assignment (Green et al., 2019), are needed to map out the range of stimulus and variability parameters that promote generalization.

Taken together, our results emphasize a role of targeted variability in achieving spatial generalization and suggest that higher stages of visual processing are involved in VPL. Variability may thus be a useful component for improving real life applications of VPL without burdening the learner with additional training time and effort.

## Declarations

### Ethics approval and consent to participate

All procedures in this study were in accordance with the principles put forward by the Declaration of Helsinki and approved by the Ethics Committee of the University Medical Center Göttingen (protocol number 29/8/17). Informed consent was obtained from all individual participants included in the study.

### Consent for publication

Consent for publication was obtained for every individual person’s data included in the study.

### Availability of data and materials

All data will be made available in a publicly accessible online repository upon acceptance of the manuscript.

### Competing interests

CMS has an editorial role with the Journal of Cognitive Enhancement. The authors have no other conflict of interest to declare.

### Funding

This project has received funding from the European Research Council (ERC) under the European Union’s Horizon 2020 research and innovation programme (Grant agreement No. 802482, to CMS). BA is supported by a scholarship from the Studienstiftung des Deutschen Volkes. CMS is supported by the German Research Foundation Emmy Noether Program (SCHW1683/2-1). The funders had no role in study design, data collection and interpretation, decision to publish, or preparation of the manuscript.

### Authors’ contributions

*Beyza Akkoyunlu:* Conceptualization, Methodology, Software, Formal analysis, Investigation, Writing - Original Draft, Visualization. *Caspar M. Schwiedrzik:* Conceptualization, Methodology, Writing - Original Draft, Supervision, Project administration, Funding acquisition.

## Acknowledgements

We would like to thank Jannik Rotzinger, Josefine-Lena Berger, and Tina Zahrie for their help in data collection, Igor Kagan for helpful discussions, and Giorgio Manenti for valuable insights during study design.

## Notes

### Competing Interest Statement

The authors have declared no competing interest.

### Summary of Updates

Following peer review in a journal, we have further expanded the results and discussion sections, respectively, on the topic of eye movements. We have also clarified statistical procedures. All results remain identical.

## References

Abrams, J., Nizam, A., & Carrasco, M. (2012). Isoeccentric locations are not equivalent: The extent of the vertical meridian asymmetry. Vision research, 52(1), 70–78. 10.1016/j.visres.2011.10.016

Adab, H. Z., & Vogels, R. (2016). Perturbation of posterior inferior temporal cortical activity impairs coarse orientation discrimination. Cerebral cortex, 26(9), 3814–3827. 10.1093/cercor/bhv178

Ahissar, E., & Arieli, A. (2012). Seeing via Miniature Eye Movements: A Dynamic Hypothesis for Vision. Frontiers in computational neuroscience, 6, 89. 10.3389/fncom.2012.00089

Ahissar, M., & Hochstein, S. (2004). The reverse hierarchy theory of visual perceptual learning. Trends in cognitive sciences, 8(10), 457–464. 10.1016/j.tics.2004.08.011

Alonso, J. M., & Martinez, L. M. (1998). Functional connectivity between simple cells and complex cells in cat striate cortex. Nature neuroscience, 1(5), 395–403. 10.1038/1609

Avidan, G., Harel, M., Hendler, T., Ben-Bashat, D., Zohary, E., & Malach, R. (2002). Contrast sensitivity in human visual areas and its relationship to object recognition. Journal of neurophysiology, 87(6), 3102–3116. 10.1152/jn.2002.87.6.3102

Bates, D., Mächler, M., Bolker, B., & Walker, S. (2015). Fitting Linear Mixed-Effects Models Using lme4. Journal of Statistical Software, 67(1). 10.18637/jss.v067.i01

Ben-Shachar, M., Lüdecke, D., & Makowski, D. (2020). effectsize: Estimation of Effect Size Indices and Standardized Parameters. Journal of Open Source Software, 5(56), 2815. 10.21105/joss.02815

Brainard, D. H. (1997). The Psychophysics Toolbox. Spatial vision, 10(4), 433–436. https://www.ncbi.nlm.nih.gov/pubmed/9176952

Cloherty, S. L., & Ibbotson, M. R. (2015). Contrast-dependent phase sensitivity in V1 but not V2 of macaque visual cortex. Journal of neurophysiology, 113(2), 434–444. 10.1152/jn.00539.2014

Cochrane, A., & Green, C. S. (2021). Assessing the functions underlying learning using by-trial and by-participant models: Evidence from two visual perceptual learning paradigms. Journal of vision, 21(13), 5. 10.1167/jov.21.13.5

Desimone, R., & Gross, C. G. (1979). Visual areas in the temporal cortex of the macaque. Brain Research, 178(2-3), 363–380. 10.1016/0006-8993(79)90699-1

Desimone, R., & Schein, S. J. (1987). Visual properties of neurons in area V4 of the macaque: sensitivity to stimulus form. Journal of neurophysiology, 57(3), 835–868. 10.1152/jn.1987.57.3.835

Diedenhofen, B., & Musch, J. (2015). cocor: A Comprehensive Solution for the Statistical Comparison of Correlations. PLoS One, 10(4), e0121945. 10.1371/journal.pone.0121945

Donovan, I., & Carrasco, M. (2018). Endogenous spatial attention during perceptual learning facilitates location transfer. Journal of vision, 18(11), 7. 10.1167/18.11.7

Dosher, B. A., & Lu, Z. L. (2005). Perceptual learning in clear displays optimizes perceptual expertise: learning the limiting process. Proceedings of the national academy of sciences of the united states of america, 102(14), 5286–5290. 10.1073/pnas.0500492102

Dosher, B. A., & Lu, Z. L. (2007). The functional form of performance improvements in perceptual learning: learning rates and transfer. Psychological science, 18(6), 531–539. 10.1111/j.1467-9280.2007.01934.x

Dosher, B. A., & Lu, Z. L. (2020). Perceptual learning. How experience shapes visual perception. MIT Press.

Douvis, S. J. (2005). Variable Practice in Learning the Forehand Drive in Tennis. Perceptual and motor skills, 101(2), 531–545. 10.2466/pms.101.2.531-545

Elkin, L. A., Kay, M., Higgins, J. J., & Wobbrock, J. O. (2021). An Aligned Rank Transform Procedure for Multifactor Contrast Tests. UIST ‘21: The 34th Annual ACM Symposium on User Interface Software and Technology, Virtual Event.

Engbert, R., & Kliegl, R. (2003). Microsaccades uncover the orientation of covert attention. Vision research, 43(9), 1035–1045. 10.1016/s0042-6989(03)00084-1

Fahle, M. (2005). Perceptual learning: specificity versus generalization. Current opinion in neurobiology, 15(2), 154–160. 10.1016/j.conb.2005.03.010

Field, D. J., & Nachmias, J. (1984). Phase reversal discrimination. Vision research, 24(4), 333–340. 10.1016/0042-6989(84)90058-0

Fiorentini, A., & Berardi, N. (1980). Perceptual learning specific for orientation and spatial frequency. Nature, 287(5777), 43–44. 10.1038/287043a0

Gattass, R., Gross, C. G., & Sandell, J. H. (1981). Visual topography of V2 in the macaque. Journal of Comparative Neurology, 201(4), 519–539. 10.1002/cne.902010405

Gladilin, E., & Eils, R. (2015). On the role of spatial phase and phase correlation in vision, illusion, and cognition. Frontiers in computational neuroscience, 9, 45. 10.3389/fncom.2015.00045

Green, C. S., Bavelier, D., Kramer, A. F., Vinogradov, S., Ansorge, U., Ball, K. K., Bingel, U., Chein, J. M., Colzato, L. S., Edwards, J. D., Facoetti, A., Gazzaley, A., Gathercole, S. E., Ghisletta, P., Gori, S., Granic, I., Hillman, C. H., Hommel, B., Jaeggi, S. M., … Witt, C. M. (2019). Improving Methodological Standards in Behavioral Interventions for Cognitive Enhancement. Journal of Cognitive Enhancement, 3(1), 2–29. 10.1007/s41465-018-0115-y

Gross, C. G., Rocha-Miranda, C. E., & Bender, D. B. (1972). Visual properties of neurons in inferotemporal cortex of the Macaque. Journal of neurophysiology, 35(1), 96–111. 10.1152/jn.1972.35.1.96

Harris, H., Gliksberg, M., & Sagi, D. (2012). Generalized perceptual learning in the absence of sensory adaptation. Current biology, 22(19), 1813–1817. 10.1016/j.cub.2012.07.059

Hennig, M. H., Kerscher, N. J., Funke, K., & Wörgötter, F. (2002). Stochastic resonance in visual cortical neurons: does the eye-tremor actually improve visual acuity? Neurocomputing, 44-46, 115–120. 10.1016/s0925-2312(02)00371-5

Higgins, J. J., & Tashtoush, S. (1994). An aligned rank transform test for interaction. Nonlinear World, 1, 201–211.

Huang, P.-C., Kingdom, F., & Hess, R. (2006). Only two phase mechanisms,+-cosine, in human vision. Vision research, 46(13), 2069–2081. 10.1016/j.visres.2005.12.020

Hubel, D. H., & Wiesel, T. N. (1962). Receptive fields, binocular interaction and functional architecture in the cat’s visual cortex. Journal of physiology, 160(1), 106–154. 10.1113/jphysiol.1962.sp006837

Hubel, D. H., & Wiesel, T. N. (1968). Receptive fields and functional architecture of monkey striate cortex. Journal of physiology, 195(1), 215–243. 10.1113/jphysiol.1968.sp008455

Hung, S. C., & Carrasco, M. (2023). Microsaccades as a long-term oculomotor correlate in visual perceptual learning. Psychonomic Bulletin & Review, 30(1), 235–249. 10.3758/s13423-022-02151-8

Kagan, I., Gur, M., & Snodderly, D. M. (2002). Spatial Organization of Receptive Fields of V1 Neurons of Alert Monkeys: Comparison With Responses to Gratings. Journal of neurophysiology, 88(5), 2557–2574. 10.1152/jn.00858.2001

Kapoula, Z., & Robinson, D. A. (1986). Saccadic undershoot is not inevitable: Saccades can be accurate. Vision research, 26(5), 735–743. 10.1016/0042-6989(86)90087-8

Karni, A., & Bertini, G. (1997). Learning perceptual skills: behavioral probes into adult cortical plasticity. Current opinion in neurobiology, 7(4), 530–535. 10.1016/s0959-4388(97)80033-5

Kassambara, A. (2023). ggpubr: ‘ggplot2’ Based Publication Ready Plots. https://CRAN.R-project.org/package=ggpubr

Kenward, M. G., & Roger, J. H. (1997). Small Sample Inference for Fixed Effects from Restricted Maximum Likelihood. Biometrics, 53(3). 10.2307/2533558

Kondat, T., Aderka, M., & Censor, N. (2023). Modulating temporal dynamics of performance across retinotopic locations enhances the generalization of perceptual learning. iScience, 26(11), 108276. 10.1016/j.isci.2023.108276

Lenth, R. V. (2023). emmeans: Estimated Marginal Means, aka Least-Squares Means. https://CRAN.R-project.org/package=emmeans

Levitt, J. B., Kiper, D. C., & Movshon, J. A. (1994). Receptive fields and functional architecture of macaque V2. Journal of neurophysiology, 71(6), 2517–2542. 10.1152/jn.1994.71.6.2517

Lu, Z.-L., & Dosher, B. A. (2004). Perceptual learning retunes the perceptual template in foveal orientation identification. Journal of vision, 4(1), 5–5. 10.1167/4.1.5

Lu, Z.-L., & Dosher, B. A. (2022). Current directions in visual perceptual learning. Nature Reviews Psychology, 1(11), 654–668. 10.1038/s44159-022-00107-2

Manenti, G. L., Dizaji, A. S., & Schwiedrzik, C. M. (2023). Variability in training unlocks generalization in visual perceptual learning through invariant representations. Current biology, 33(5), 817–826 e813. 10.1016/j.cub.2023.01.011

Maniglia, M., & Seitz, A. R. (2018). Towards a whole brain model of Perceptual Learning. Current Opinion in Behavioral Sciences, 20, 47–55. 10.1016/j.cobeha.2017.10.004

Melloni, L., Schwiedrzik, C. M., Rodriguez, E., & Singer, W. (2009). (Micro)Saccades, corollary activity and cortical oscillations. Trends in cognitive sciences, 13(6), 239–245. 10.1016/j.tics.2009.03.007

Otto, T. U., Herzog, M. H., Fahle, M., & Zhaoping, L. (2006). Perceptual learning with spatial uncertainties. Vision research, 46(19), 3223–3233. 10.1016/j.visres.2006.03.021

Perry, L. K., Samuelson, L. K., Malloy, L. M., & Schiffer, R. N. (2010). Learn Locally, Think Globally: Exemplar Variability Supports Higher-Order Generalization and Word Learning. Psychological science, 21(12), 1894–1902. 10.1177/0956797610389189

Raviv, L., Lupyan, G., & Green, S. C. (2022). How variability shapes learning and generalization. Trends in cognitive sciences, 26(6), 462–483. 10.1016/j.tics.2022.03.007

Ringach, D. L. (2002). Spatial structure and symmetry of simple-cell receptive fields in macaque primary visual cortex. Journal of neurophysiology, 88(1), 455–463. 10.1152/jn.2002.88.1.455

Rucci, M., Iovin, R., Poletti, M., & Santini, F. (2007). Miniature eye movements enhance fine spatial detail. Nature, 447(7146), 851–854. 10.1038/nature05866

Schiller, P. H., Finlay, B. L., & Volman, S. F. (1976). Quantitative studies of single-cell properties in monkey striate cortex. I. Spatiotemporal organization of receptive fields. Journal of neurophysiology, 39(6), 1288–1319. 10.1152/jn.1976.39.6.1288

Schoups, A. A., Vogels, R., & Orban, G. A. (1995). Human perceptual learning in identifying the oblique orientation: retinotopy, orientation specificity and monocularity. Journal of physiology, 483 (Pt 3), 797–810. 10.1113/jphysiol.1995.sp020623

Schoups, A. A., Vogels, R., Qian, N., & Orban, G. A. (2001). Practising orientation identification improves orientation coding in V1 neurons. Nature, 412(6846), 549–553. 10.1038/35087601

Serre, T., & Riesenhuber, M. (2004). Realistic Modeling of Simple and Complex Cell Tuning in the HMAX Model, and Implications for Invariant Object Recognition in Cortex. CSAIL Technical Reports, 052. http://hdl.handle.net/1721.1/30491

Shapley, R. (1985). The importance of contrast in the perception of brightness and form. Transactions of the American Philosophical Society, 75(6), 20–29.

Shiu, L. P., & Pashler, H. (1992). Improvement in line orientation discrimination is retinally local but dependent on cognitive set. Perception & psychophysics, 52(5), 582–588. 10.3758/bf03206720

Sowden, P. T., Rose, D., & Davies, I. R. (2002). Perceptual learning of luminance contrast detection: specific for spatial frequency and retinal location but not orientation. Vision research, 42(10), 1249–1258. 10.1016/s0042-6989(02)00019-6

Talluri, B. C., Hung, S.-C., Seitz, A. R., & Seriès, P. (2015). Confidence-based integrated reweighting model of task-difficulty explains location-based specificity in perceptual learning. Journal of vision, 15(10), 17. 10.1167/15.10.17

Tan, Q., Wang, Z., Sasaki, Y., & Watanabe, T. (2019). Category-induced transfer of visual perceptual learning. Current biology, 29(8), 1374–1378. 10.1016/j.cub.2019.03.003

Victor, J. D., & Purpura, K. P. (1998). Spatial phase and the temporal structure of the response to gratings in V1. Journal of neurophysiology, 80(2), 554–571. 10.1152/jn.1998.80.2.554

Vogels, R., & Orban, G. A. (1994). Activity of inferior temporal neurons during orientation discrimination with successively presented gratings. Journal of neurophysiology, 71(4), 1428–1451. https://www.ncbi.nlm.nih.gov/pubmed/8035226

Vukatana, E., Graham, S. A., Curtin, S., & Zepeda, M. S. (2015). One is Not Enough: Multiple Exemplars Facilitate Infants’ Generalizations of Novel Properties. Infancy, 20(5), 548–575. 10.1111/infa.12092

Watanabe, T., & Sasaki, Y. (2015). Perceptual Learning: Toward a Comprehensive Theory. Annual Review of Psychology, 66(1), 197–221. 10.1146/annurev-psych-010814-015214

Wickham, H. (2016). ggplot2: Elegant Graphics for Data Analysis. Springer. 10.1007/978-3-319-24277-4

Wilson, J. R., & Sherman, S. M. (1976). Receptive-field characteristics of neurons in cat striate cortex: Changes with visual field eccentricity. Journal of neurophysiology, 39(3), 512–533. 10.1152/jn.1976.39.3.512

Wobbrock, J. O., Findlater, L., Gergle, D., & Higgins, J. J. (2011). The Aligned Rank Transform for Nonparametric Factorial Analyses Using Only Anova Procedures. CHI ‘11: Proceedings of the SIGCHI Conference on Human Factors in Computing Systems, 143–146. 10.1145/1978942.1978963

Xiao, L. Q., Zhang, J. Y., Wang, R., Klein, S. A., Levi, D. M., & Yu, C. (2008). Complete transfer of perceptual learning across retinal locations enabled by double training. Current biology, 18(24), 1922–1926. 10.1016/j.cub.2008.10.030

Zaman, J., Chalkia, A., Zenses, A.-K., Bilgin, A. S., Beckers, T., Vervliet, B., & Boddez, Y. (2021). Perceptual variability: Implications for learning and generalization. Psychonomic Bulletin & Review, 28(1), 1–19. 10.3758/s13423-020-01780-1

Zhang, T., Xiao, L. Q., Klein, S. A., Levi, D. M., & Yu, C. (2010). Decoupling location specificity from perceptual learning of orientation discrimination. Vision research, 50(4), 368–374. 10.1016/j.visres.2009.08.024

Zhaoping, L., Herzog, M. H., & Dayan, P. (2003). Nonlinear ideal observation and recurrent preprocessing in perceptual learning. Network, 14(2), 233–247. https://www.ncbi.nlm.nih.gov/pubmed/12790183

