## Supplementary Material for "Task-irrelevant phase but not contrast variability unlocks generalization in visual perceptual learning"

**Supplementary information**

*Learning curves*

To investigate the functional form of learning, we fitted nonlinear functions, a 3-parameter exponential ($start + (asymptote - start)^{time \times rate}$) and a 3-parameter power function ($start + (asymptote - start) \times time^{rate}$) using the TEfits package (Cochrane, 2020) to each participant's thresholds at the respective training locations (3-pre learning, 20 training, 3-post learning blocks). We chose these two functions as they have both been suggested to capture the dynamics of VPL (Dosher & Lu, 2005, 2007). To quantify the fit, we calculated Bayesian Information Criteria (BIC) and compared these BIC values statistically using ANOVA. Power function fits resulted in lower BIC values (power law mean BIC -21.80, SEM 4.95; exponential function mean BIC -23.57, SEM 4.94); this difference was statistically significant ($F(1,25)=6.43$, $p=0.02$, $\eta_p^2=0.2$). These findings are in line with previous findings by Dosher and Lu (2005), while other studies report a better fit with an exponential rather than a power function (Cochrane & Green, 2021; Dosher & Lu, 2007). It is possible that methodological differences explain these discrepant results. For example, we as well as Dosher and Lu (2005) trained with small, localized stimuli, while Dosher and Lu (2007) and Cochrane & Green (2021) used full-field displays. This may have lead to different difficulty levels of the respective tasks, which may have resulted in different learning curves (Cochrane & Green, 2021).

BIC values did not change depending on randomization or training location nor the interactions of those factors (randomization $F(1,25)=0.59$, $p=0.44$, $\eta_p^2=0.02$; training location $F(1,25)=2.81$, $p=0.10$, $\eta_p^2=0.1$; function × randomization $F(1,25)=0.23$, $p=0.62$, $\eta_p^2=9.12\times10^{-3}$; function × training location $F(1,25)=2.78$, $p=0.10$, $\eta_p^2=0.1$; randomization × training location $F(1,25)=0.66$, $p=0.42$, $\eta_p^2=0.03$; function × randomization × training location $F(1,25)=2.19$, $p=0.15$, $\eta_p^2=0.08$). Hence, all experimental groups showed similar learning curves over sessions.

After establishing the functional form of learning, we also tested the power law function parameters statistically, and found that none of the three parameters changed depending on randomization or training location (Rate: randomization $F(1,25)=0.32$, $p=0.57$, $\eta_p^2=0.01$; training location $F(1,25)=0.22$, $p=0.64$, $\eta_p^2= 8.72\times10^{-3}$; randomization × training location $F(1,25)=2.06$, $p=0.16$, $\eta_p^2=0.08$; Start: randomization $F(1,25)=0.26$, $p=0.60$, $\eta_p^2=0.01$; training location $F(1,25)=0.48$, $p=0.49$, $\eta_p^2=0.02$, randomization × training location $F(1,25)=1.21$, $p=0.28$, $\eta_p^2=0.05$; Asymptote: randomization

F(1,25)=0.40, p=0.53, $\eta_p^2$=0.02; training location F(1,25)=1.90, p=0.18, $\eta_p^2$=0.07; randomization × training location F(1,25)=0.11, p=0.73, $\eta_p^2$=4.38×10$^{-3}$). In sum, all experimental groups show characteristics of VPL by improving their performance gradually over multiple sessions, and moreover we do not find any evidence for different learning processes across groups.

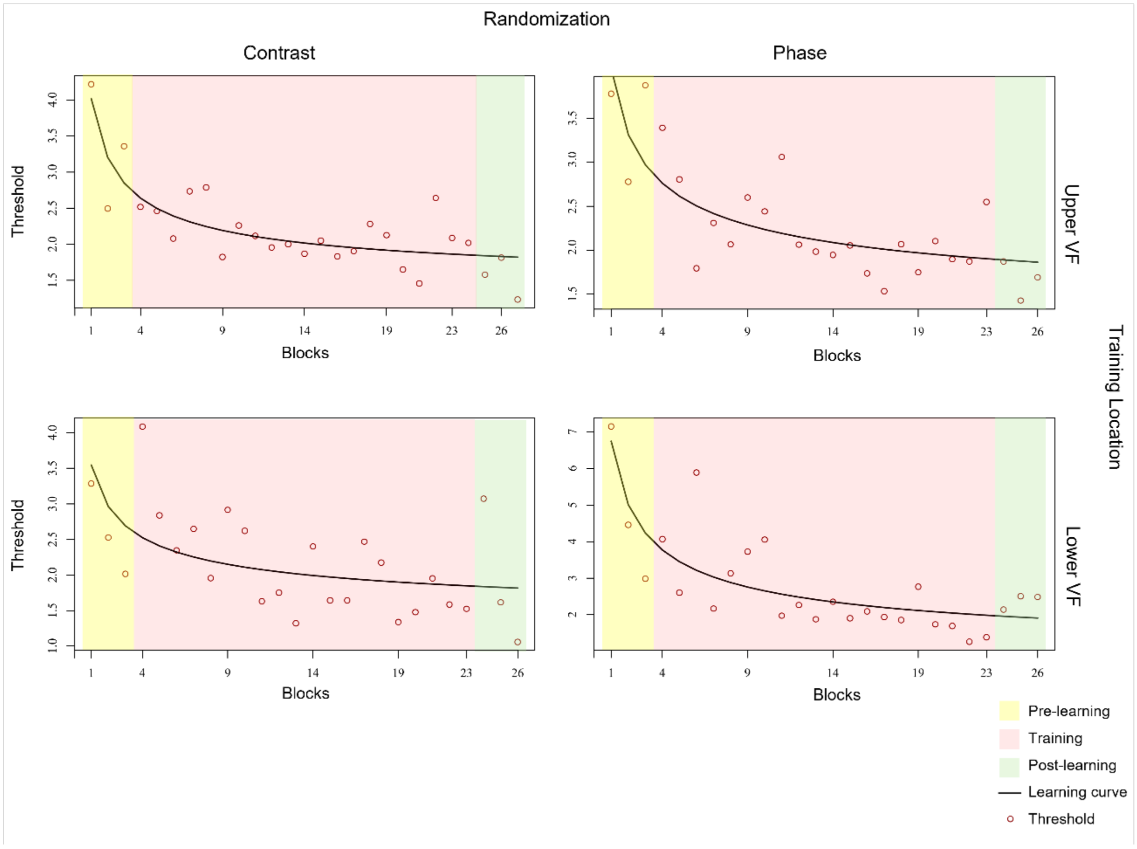


**Supplementary Figure S1.** Example learning fits from each group. Black continuous lines represent power law function fits to the thresholds (red circles). Yellow background represents thresholds from pre-learning measurements, red background training blocks, and green background shows thresholds from post-learning measurements.

Supplementary Table 1. rmANOVA results for the fixed effects in the linear mixed effects model

| Term | F | Df | Df.res | Pr(>F) |
|---|---|---|---|---|
| Randomization | 0.749 | 1 | 24.991 | 0.394 |
| Time | 361.319 | 1 | 469.029 | 0.000 |
| Location | 25.172 | 2 | 469.028 | 0.000 |
| Training Location | 3.710 | 1 | 24.991 | 0.065 |
| Randomization:Time | 6.703 | 1 | 469.031 | 0.009 |
| Randomization:Location | 2.583 | 2 | 469.029 | 0.076 |
| Time:Location | 0.717 | 2 | 469.031 | 0.488 |
| Randomization:Training Location | 1.099 | 1 | 24.991 | 0.304 |
| Time:Training Location | 3.942 | 1 | 469.031 | 0.047 |
| Location:Training Location | 10.644 | 2 | 469.028 | 0.000 |
| Randomization:Time:Location | 3.317 | 2 | 469.031 | 0.037 |
| Randomization:Time:Training Location | 0.219 | 1 | 469.031 | 0.639 |
| Randomization:Location:Training Location | 6.002 | 2 | 469.029 | 0.002 |
| Time:Location:Training Location | 0.120 | 2 | 469.031 | 0.886 |
| Randomization:Time:Location:Training Location | 3.407 | 2 | 469.031 | 0.033 |

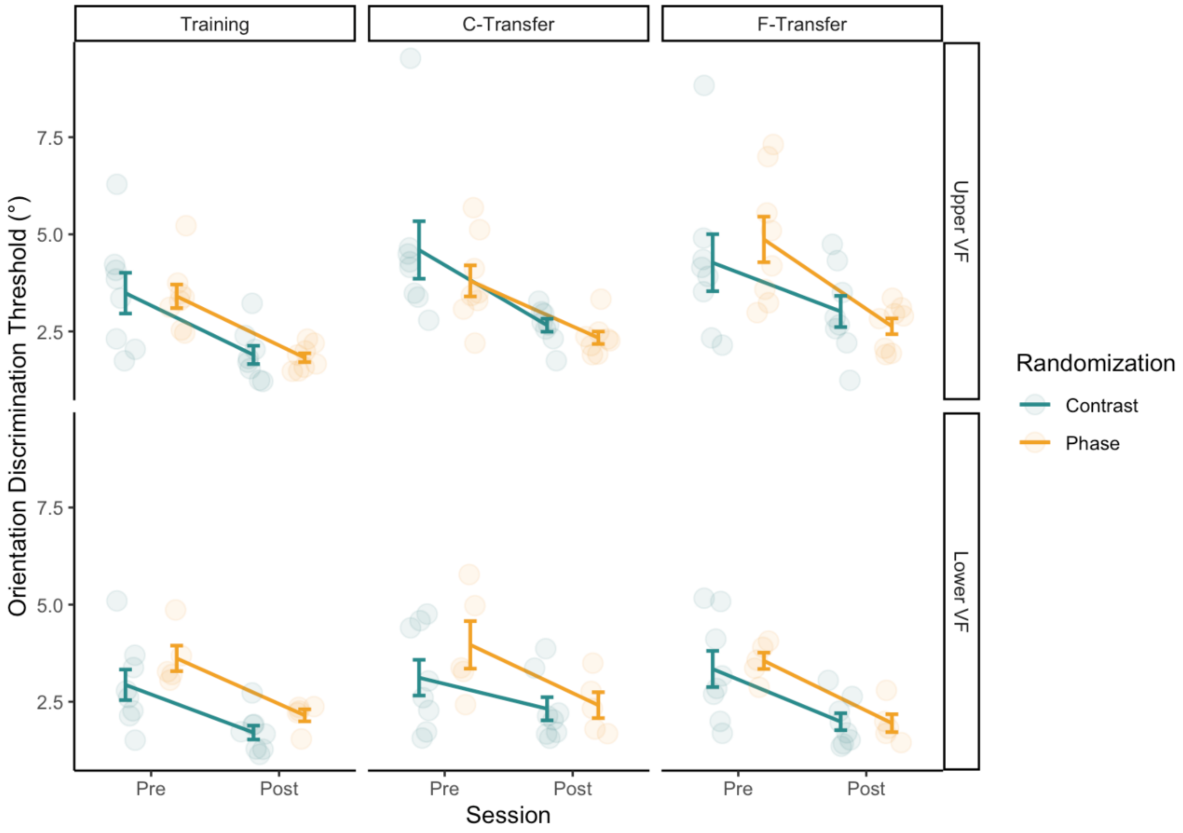


**Supplementary Figure S2.** Orientation discrimination thresholds of all experimental groups at different visual field locations and time points. The upper section of the figure shows the experimental groups which were trained in upper visual field, the lower section shows groups trained in lower visual field. While there was a statistically significant four-way interaction between randomization, time, location, and training location, the trends reported for the two transfer locations are largely consistent across the visual field. Specifically, despite the significant four-way interaction, post hoc tests yielded no statistically significant differences between the differential learning effects on thresholds in the contrast versus phase group for subjects trained in the upper versus lower visual field (Far Transfer location t(469)=-0.985, p=0.4874, Cohen's d=0.09; Close Transfer location t(469)=1.399, p=0.4873, Cohen's d=0.13; Training location t(469)=-0.413, p=0.6797, Cohen's d=0.04, corrected for multiple comparisons using the False Discovery Rate, but uncorrected p-values were also not statistically significant). Hence, the difference in generalization does not seem to depend on the training location. Each dot represents the mean of a subject's threshold at the respective location, error bars represent the standard error of the mean.

### Supplementary references


Cochrane, A. (2020). TEfits: Nonlinear regression for time-evolving indices. *Journal of Open Source Software*, *5*(52), 2535. https://doi.org/10.21105/joss.02535

Cochrane, A., & Green, C. S. (2021). Assessing the functions underlying learning using by-trial and by-participant models: Evidence from two visual perceptual learning paradigms. *Journal of vision*, *21*(13), 5. https://doi.org/10.1167/jov.21.13.5

Dosher, B. A., & Lu, Z. L. (2005). Perceptual learning in clear displays optimizes perceptual expertise: learning the limiting process. *Proceedings of the national academy of sciences of the united states of america*, *102*(14), 5286-5290. https://doi.org/10.1073/pnas.0500492102

Dosher, B. A., & Lu, Z. L. (2007). The functional form of performance improvements in perceptual learning: learning rates and transfer. *Psychological science*, *18*(6), 531-539. https://doi.org/10.1111/j.1467-9280.2007.01934.x
